# Transfer of graded information through gated receptivity to widely broadcast signals

**DOI:** 10.1101/2025.09.30.679638

**Authors:** Lindsey S. Brown, NaYoung So, L. F. Abbott, Michael N. Shadlen, Mark S. Goldman

## Abstract

Making accurate decisions requires the brain to maintain evolving representations of the evidence supporting different possible choices. The population of neurons that maintains this evolving representation may change over the course of the decision-making process. Such changes may result from a switch in the set of neurons representing the relevant behavioral output or a switch in the set of neurons receiving task-relevant information. Recent work has shown how intervening actions like eye movements or navigation to a new location can shift the set of neurons that encode subsequent inputs and outputs. Here, we present a computational model of how analog-valued information that supports an evolving decision can be flexibly transferred between populations of neurons without changing the synaptic connectivity of the underlying network. This is accomplished through a dynamic gating mechanism in which information is widely broadcast throughout a network, but only those neurons receiving a gating signal are receptive to this information. Using this framework, we provide a mechanistic explanation for recent experimental results in which information supporting an evolving decision was shown to be transferred between different populations of parietal cortex neurons during intervening smooth pursuit and saccadic eye movements. This mechanism enables organisms to maintain a continuous decision process despite changing frames of reference, offering a potentially general framework for cognitive continuity in dynamic environments.

## Introduction

Many decisions require gathering multiple pieces of information that are available at different times and locations. For example, suppose a person is faced with a decision that involves accumulating evidence for two competing choices and then pressing one of two buttons, based on which choice has more evidence supporting it. Previous work^1–5^ suggests that the neurons in the brain representing the evidence for each choice are explicitly tied to the final movement required to report the decision (i.e., to the location of the buttons in this example). Thus, if the observer moves their body while accumulating evidence for the decision, then this action will shift both the required movement and the set of neurons that encodes the decision. Despite such reorienting actions, the brain must maintain a continuous representation of the evolving decision.

Several studies of decision-making have considered tasks in which, due to intervening actions by an animal, the information relevant to a decision is transferred from one population of neurons to a different population. Rodent studies have observed that information about the accumulated evidence for a decision is transferred sequentially between neural populations while an animal navigates^6–10^. In nonhuman primates, a recent study that we focus upon here explored decision making in the face of intervening eye movements^11^. Monkeys were required to determine the net direction of motion in random dot motion stimuli presented in two brief pulses. The two pulses were separated by a set of intervening eye movements that changed the gaze direction (Fig. 1A,B), either gradually in smooth pursuit movements or abruptly in saccadic eye movements. The monkeys integrated motion inputs from before and after the gaze shift in each trial. Neural recordings revealed that the parietal neurons representing the accumulated evidence from the first motion pulse became uninformative after the eye movement, as their response field no longer contained the choice target. However, before it was lost, the accumulated evidence for the decision was transferred to a different population of neurons whose response fields aligned with the choice target relative to the new gaze direction. This new population resumed the decision process, updating the previously accumulated evidence with new motion evidence.

**Figure 1.**
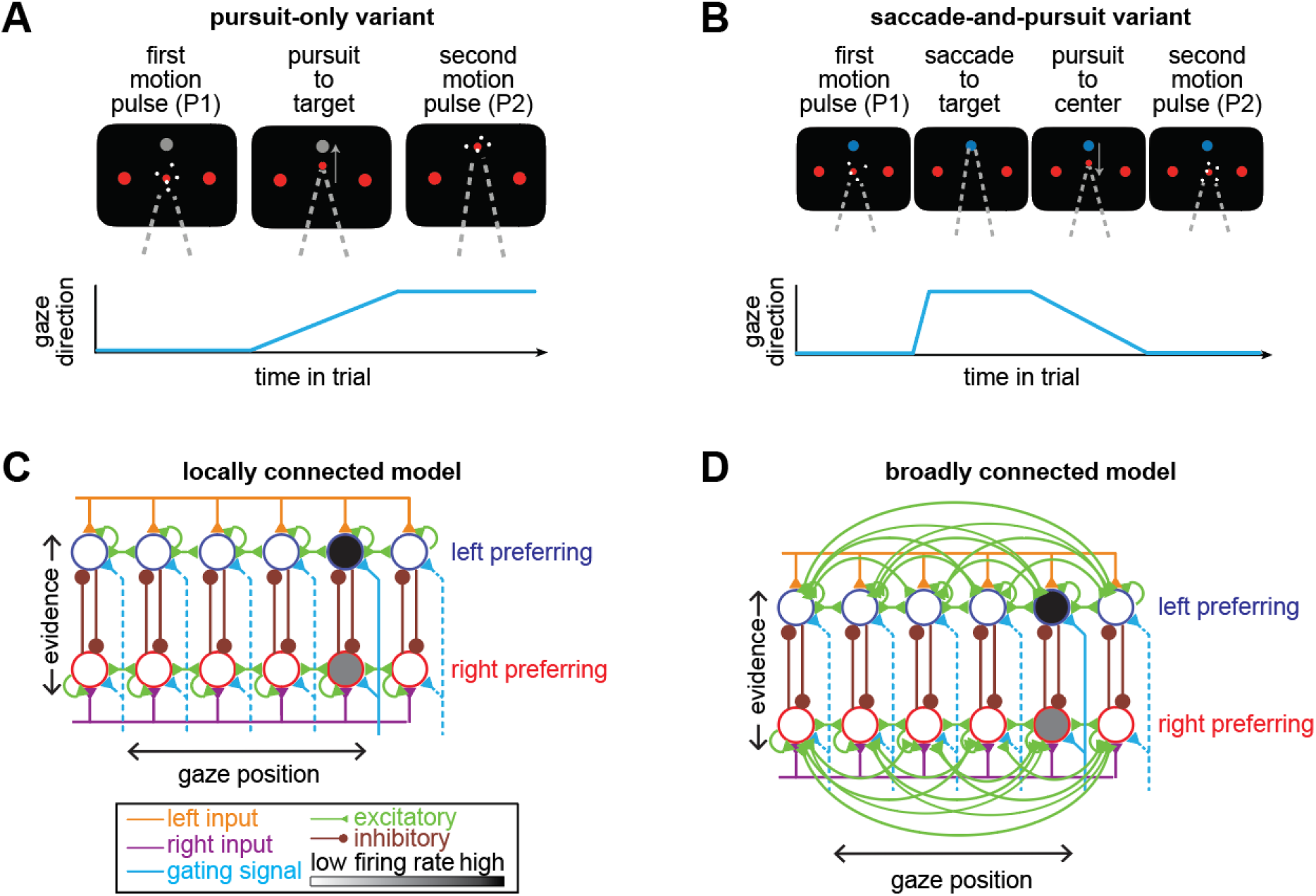
Variants of the two-pulse random dot motion discrimination task and candidate models of evidence accumulation, maintenance, and transfer. (A) *Top*: Sequence of the trial in the pursuit-only variant of the two-pulse task. Two brief motion pulses are shown, one before and one after the intervening pursuit eye movement, which brings the monkey’s gaze direction from the initial fixation point at the center to the gray target. *Bottom*: Schematic of the monkey’s gaze direction during different stages of the task (cyan). (B) Same as A but for the saccade-and-pursuit variant of the two-pulse task, in which the monkey needs to make a saccade to the blue target after the first motion pulse and a smooth pursuit eye movement back to the initial fixation point at the center. (C) Schematic of the locally connected model. Neurons (circles) have local excitatory (green) connections with adjacent neurons in the population with the same side preference (left-preferring (blue circles) and right-preferring (red circles)) and mutually inhibitory (brown) connections between neurons with opposite side preference. All neurons at a given gaze position receive the same gating signal (cyan). External inputs conveying leftward (orange) or rightward (purple) motion project to all neurons of the corresponding side preference. (D) Same as C but for the broadly connected model, in which neurons have excitatory connections with all other neurons that share the same side preference. Panels A and B are adapted from So & Shadlen (2022)^11^. Panel C is adapted from Brown et al. (2026)^12^.

This finding provides a striking example of flexible information transfer between neurons. It enables a neural population that encodes decisions in an eye-centered (oculocentric) reference frame to support representations that are invariant to gaze shifts. Similar information transfer may underlie continuity in other domains, such as spatial navigation, where internal representations of a navigational goal must be maintained across movement through space^6–10^. However, eye movements, especially saccades, pose unique challenges due to their abrupt and rapid nature. While smooth pursuit eye movements change the gaze direction continuously, similar to movements in navigation, saccadic eye movements involve rapid shifts between discontiguous gaze directions, demanding robust and flexible mechanisms to transfer graded information between distinct populations of neurons.

Here, we present a circuit model that accounts for the robust and flexible transfer of graded evidence across both smooth pursuit and saccadic eye movements, as observed in So & Shadlen (2022)^11^. We first consider two possible models, one with local connectivity between neurons that have neighboring response fields^12^ and a second with broad connectivity in which signals are widely broadcast throughout the network. Both models operate without changes in synaptic connectivity; instead, they rely on a dynamic gating mechanism that shifts the receptivity of downstream neurons to the graded evidence signals. We then analyze the temporal and spatial characteristics of the transfer in the data. The rapid and saltatory transfer across saccades is better captured by the model with broad connectivity. Finally, we discuss how the proposed mechanism of gating access to widely broadcast signals enables continuous integration, maintenance, and transfer of evidence across shifting frames of reference, preserving and extending the fidelity of cognitive processes such as decision-making, memory, and attention under dynamic behavioral conditions.

## Results

### Models of accumulation, maintenance, and transfer of graded information across eye movements

Single neurons in LIP are known to represent the accumulation of noisy evidence that supports the decision to choose a target in the neuron’s response field. However, these neurons only maintain this graded evidence representation while the choice target lies in their response field. When the monkey shifts its gaze, the choice target will lie in the response field of a different population of LIP neurons, and the representation is transferred to this population^11^. This transfer of information supports the continuous representation of accumulated evidence across the intervening eye movements both when the gaze shift is smooth (as in smooth pursuit; Fig. 1A) and when it is abrupt (as in a saccade; Fig. 1B). Thus, neurons in LIP are part of a circuit that performs two key computations, the accumulation of evidence at any fixed gaze direction and the transfer of this graded evidence between populations encoding different gaze directions. In what follows, we develop a model of this process, focusing primarily on the less studied operation of graded information transfer.

The first computation – accumulating and maintaining evidence – has been well studied both experimentally and computationally. Many experimental observations^13–18^ have been compactly described within the framework of drift-diffusion models^19–21^ and such models have been mechanistically realized in neural circuit models based on recurrent feedback among populations of neurons^22–26^. Aligned with the predictions of these models, the activity in LIP at a fixed gaze direction is well described by such drift-diffusion processes^1,27–30^ and their network realizations^31–33^. In our model, we assume that at each fixed gaze direction, there are two subpopulations of neurons that represent accumulated evidence for each one of the choice targets, through a balance of excitation within the same-choice-preferring population and mutual inhibition between the populations with opposite-choice preferences.

The second computation – the transfer of graded information between populations – is less commonly studied, although it has been considered in the case of a navigation task in which accumulated evidence must be maintained as information is passed between neurons encoding different spatial positions in a maze^7,10,12^. In the case of accumulation of evidence in the presence of gaze shifts, an eye movement can take the gaze from any initial to any final direction. Hence, the current neural population maintaining the activity must be able to send its accumulated evidence representation to neurons with response fields corresponding to all other gaze directions (i.e., those neurons that, at a given gaze direction, have response fields that contain the choice target). Specifically, the representation must be able to be transferred between the current population of neurons with response fields that contain the choice target when the gaze is at its initial direction to any final population of neurons with response fields that contain the choice target when the gaze is at its final direction. Within LIP, this transfer could be done in two manners: 1) a sequential, chain-like transfer of activity across neurons with adjacent response fields, starting from the current population and ending with the final population, or 2) a single saltatory transfer, jumping directly from the current to the final population. In this second case, each neural population must directly connect to all other populations in order to be able to directly transfer information between any initial and final gaze direction. This implies that the sending population globally broadcasts its information, so the transfer of information must be controlled by changing the receptivity of the receiving neurons^6^. Here, we formalize a proposed mechanism underlying such flexible transfer: information is widely broadcast by the sending neurons, while a gating signal brings the neural activity of the receiving neurons above the firing threshold to make these neurons receptive to the widely broadcast signals.

We instantiate these two means of transferring activity through two models with different connectivity schemes. In the first model, which we call the “locally connected” model, neurons form two locally connected chains, with each location along the chain representing one gaze direction along a one-dimensional eye movement trajectory in the two-dimensional visual space (Fig. 1C), similar to a previous model of navigation-based decision-making tasks^12^. In the second model, which we call the “broadly connected” model, neurons are broadly connected within the population of the same choice preference in an all-to-all manner (Fig. 1D).

In the locally connected model, the firing rate *r*_*i,L*_ of the *i*^th^ neuron in the left-preferring population is given by

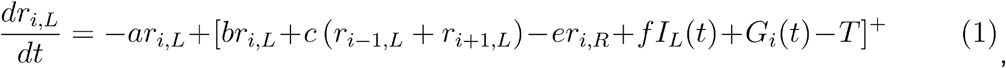

where *i* indexes the response field location and []^+^ denotes rectification (i.e., setting negative values to zero). In the broadly connected model, the evolution of the firing rate is similar, but each neuron receives input broadly from all other neurons in the same-choice-preferring population,

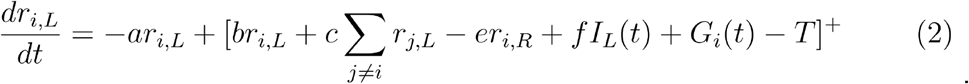

The firing rate *r*_*i,R*_ of the *i*^th^ neuron in the right-preferring population in each model is defined analogously by interchanging *r*_*i,L*_ and *r*_*i,R*_ for all *i* and substituting *I*_*R*_ for *I*_*L*_. In both models, the activity of the neuron decays exponentially in the absence of input at rate *a*. Each neuron excites itself with weight *b* and excites other neurons within the same population that it is connected to with strength *c* (Fig. 1C,D, green connections). Each neuron inhibits the neuron with a corresponding response field in the opposing population with strength *e* (Fig. 1C,D, brown connections). Evidence inputs to the model are given by *I*_*L*_(*t*) *=* |*C*|***1***_*L*_(*t*), where *C* is the signed coherence of the random dot motion, and ***1***_***L***_(*t*) *=* 1 if the motion is on and to the left (and correspondingly, ***1***_***R***_(*t*) *=* 1 if the motion is on and to the right) and 0 otherwise. *G*_*i*_(*t*) is the gating signal that brings the neuron above its firing threshold *T* when the choice targets lie within the neuron’s response field. Note that each model neuron should not be thought of as a single biological neuron but as the average response of a population of neurons with similar spatial tuning.

The gating signal ensures that only neurons with the choice target in their response field actively accumulate evidence and maintain this graded information. At a fixed gaze direction, both models reduce to traditional neural instantiations of the drift-diffusion model, so the same tuning conditions for integration apply^24^. Additionally, the connections within the same-choice-preferring population *c* must be equal to the decay rates *a* for the graded information to be preserved (neither gained nor lost) when transferred across gaze directions (see Methods). Neither model requires fine-tuning of the gating signal. Instead, the gating signal only needs to 1) target the appropriate neurons that have a response field containing the choice target and 2) be above the firing threshold *T* to make the neuron receptive to its other inputs.

We simulated these models on the two variants of the two-pulse, random dot motion discrimination task^11^ (Fig. 1A,B) to test under what conditions these models achieve the transfer of accumulated evidence between populations when a saccade or pursuit eye movement changes the population of neurons with response fields that contain one of the choice targets.

### Smooth transfer of graded information during smooth pursuit eye movement

In the pursuit-only variant of the task, graded decision-related information is transferred between LIP neurons during the smooth pursuit eye movement (Fig. 1A)^11^. Before the pursuit eye movement, the monkey accumulates evidence based on the first motion pulse (P1), and a population of neurons whose response fields are aligned with one of the choice targets shows ramping activity, with the slope reflecting the evidence for the decision (i.e., the motion strength and direction of P1) (Fig. 2A,B, top left). After the pulse, the ramping in these neurons stops, but the activity maintains a graded level reflecting the provisional decision until the onset of the pursuit eye movement. As the monkey’s gaze direction shifts during the pursuit, the decision-related activity shifts from this initially responsive population (population 1) of neurons to a different set of neurons (population 2) whose response fields now encompass the choice target (Fig. 2A, bottom middle). This transfer is smooth, with activity propagating through neurons that sequentially acquire the choice target in their response fields over the ∼750 ms duration of smooth pursuit (Fig. 2B, bottom middle). After the pursuit, only those neurons with response fields that contain the choice target at the endpoint of the pursuit eye movement update their representation of the decision by integrating new evidence from the second motion pulse (P2), building on the initial graded state so that the final activity reflects both pulses (Fig. 2A,B, bottom right).

**Figure 2.**
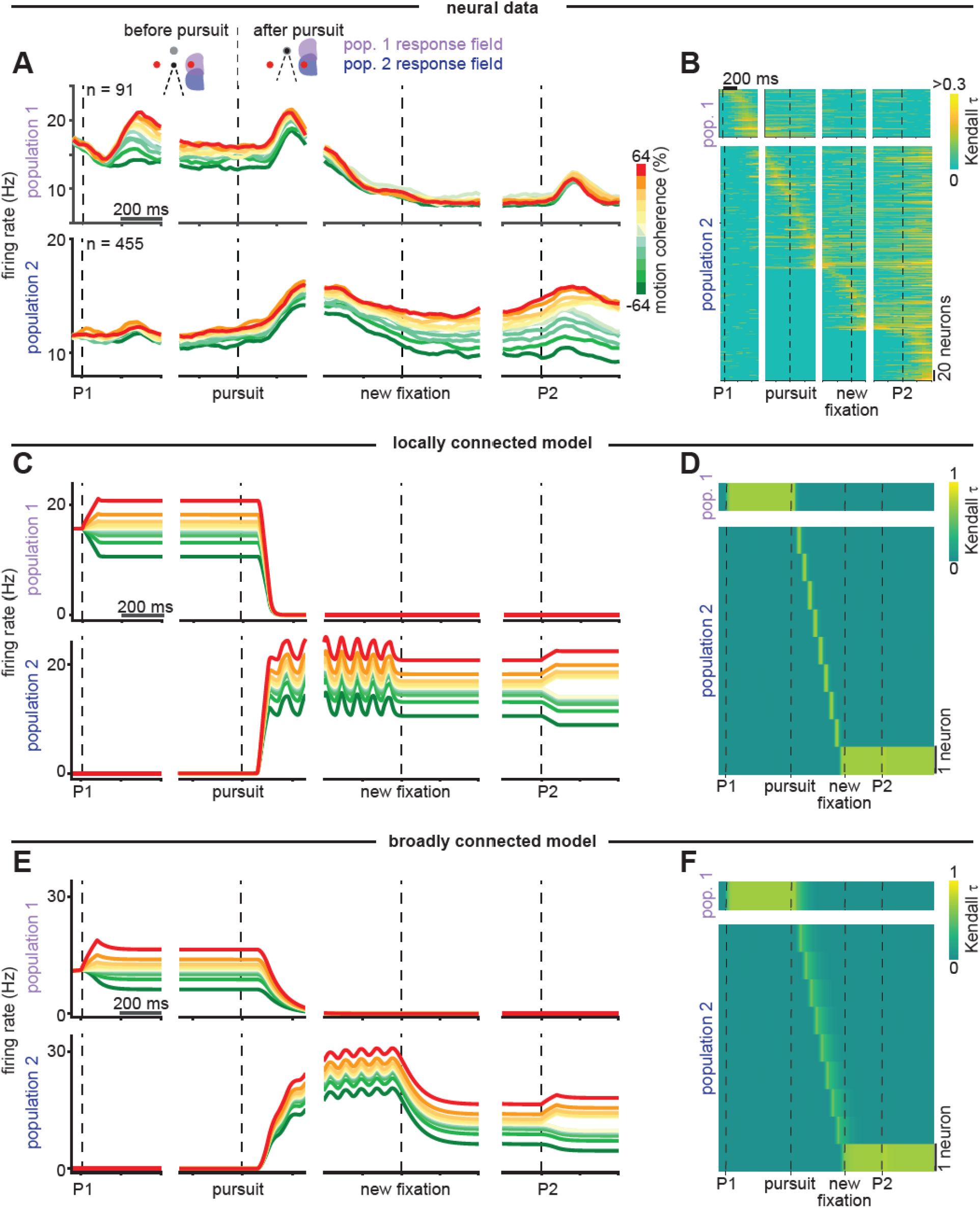
Smooth transfer of graded information during smooth pursuit eye movement. (A) Average activity of the LIP neurons that represent the decision before the pursuit eye movement (population 1, *top*), and during and after the pursuit eye movement (population 2, *bottom*). Colors indicate motion strength and direction. (B) Decision-related activity of individual neurons (rows). The heatmap displays the strength of correlation (Kendall *τ*) of the activity with the motion strength of the first pulse P1. The neurons are ordered by the time of peak correlation. (C) As in A but for the sum total of the trial-averaged activity of simulated neurons from the locally connected model. (D) Same as B but for simulated neurons from the locally connected model. Oscillations in population 2 activity during the pursuit eye movement reflect the discrete, sequential gating on and off of individual neurons in the model. (E, F) Same as C, D but for simulated neurons from the broadly connected model. Panels A and B are adapted from So & Shadlen (2022)^11^, with population 2 expanded to include neurons exhibiting decision-related activity only during the pursuit eye movement.

Simulations of both the locally and broadly connected models on this task reproduced the integration and transfer of graded information across neurons (Fig. 2C-F). The transfer of the graded decision information is also apparent at the level of individual neurons in both models (Fig. 2D,F), as in the neural recordings. During the smooth pursuit eye movement, the information was smoothly transferred across neurons corresponding to different gaze directions. Overall, it is not surprising that both models capture this smooth transfer of the graded information, since both the locally and broadly connected models contain the local connections that mediate the transfer of information between neurons with neighboring response fields. Next, we show that the predictions of these models differ strongly for the abrupt changes in gaze direction that occur across a saccade.

### Saltatory transfer of graded information during saccadic eye movement

In the saccade-and-pursuit variant of the task (Fig. 1B), the graded decision information is transferred rapidly across LIP neurons as gaze direction shifts abruptly during a saccade. As in the pursuit-only variant, a graded representation of the decision information emerges in the initially responsive neurons during the first motion pulse P1 (population 1; Fig. 3A,B, top left). Then, instead of making a smooth pursuit, the monkey makes a saccade, and the graded decision information represented in these neurons is rapidly transferred to those neurons whose response fields now align with the choice target (population 2; Fig. 3A,B, bottom middle). Before the next motion pulse P2, the monkey returned its gaze to the center via a smooth pursuit eye movement. As the gaze returns, the decision information is transferred back to the population 1 neurons, which update their representation with the new evidence P2 (Fig. 3A,B, top right).

**Figure 3.**
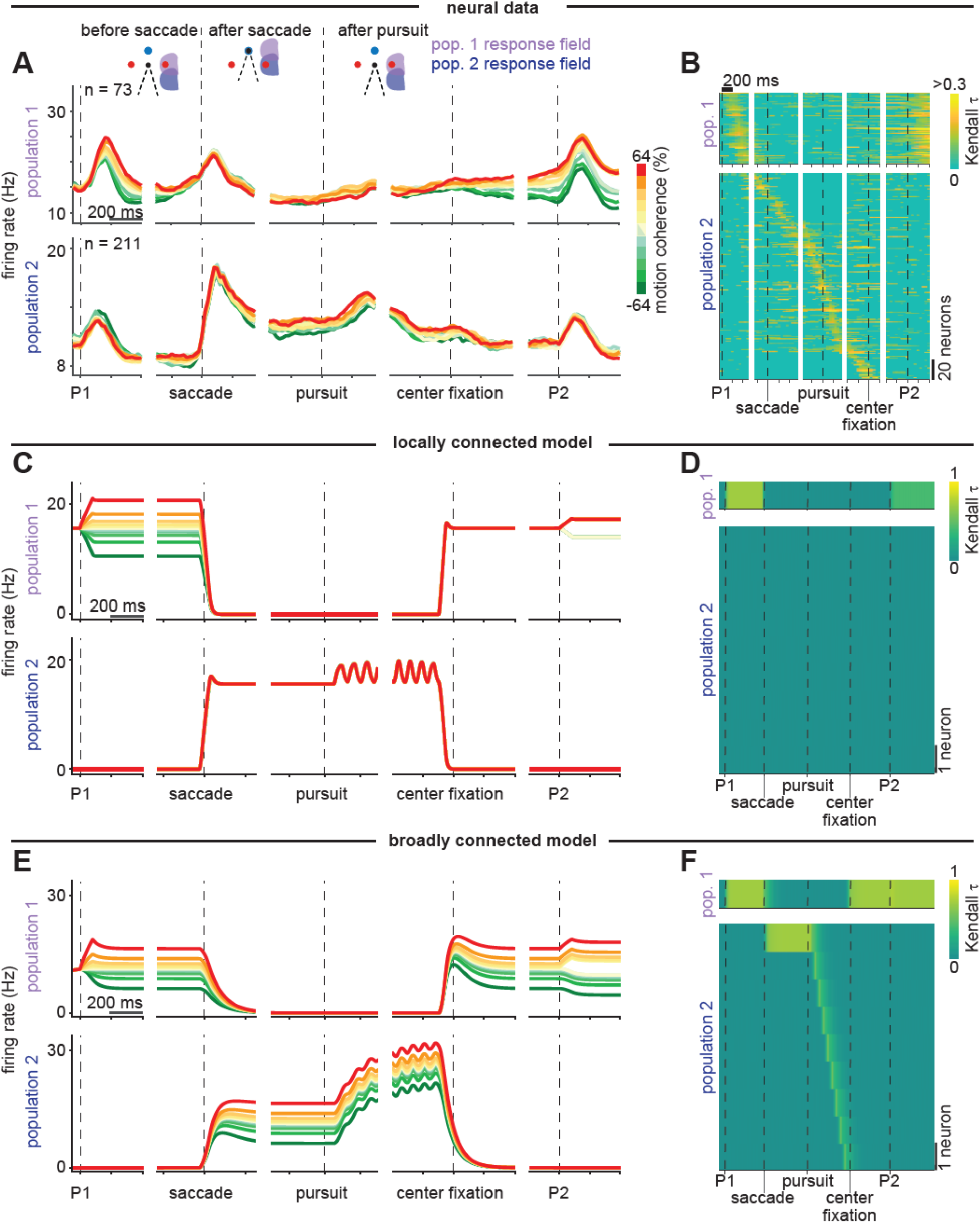
Saltatory transfer of graded information during saccadic eye movement. (A) Average activity of the LIP neurons during the saccade-and-pursuit variant of the two-pulse task (*top*: population 1, *bottom*: population 2). Colors indicate motion strength and direction. (B) Decision-related activity of individual neurons (rows), represented as the strength of correlation of the activity with motion strength of the first pulse P1 (Kendall *τ*). The neurons are ordered by the time of peak correlation. (C) Same as A, but for the sum total of the trial-averaged activity of the simulated neurons from the locally connected model. Oscillations in population 2 activity during the pursuit eye movement reflect the discrete, sequential gating on and off of individual neurons in the model. (D) Same as B but for simulated neurons from the locally connected model. (E, F) Same as C, D but for simulated neurons from the broadly connected model. Panels A and B are adapted from So & Shadlen (2022)^11^.

The two models differ in their ability to transfer graded decision information across a saccade. The locally connected model fails to transfer the graded information; instead, the simulated activity in the population 2 neurons reflects only the gating signal, indicating whether the neurons have the choice target in their response fields (Fig. 3C,D). This failure occurs because the gating signal advances between neighboring neurons over a time interval shorter than the time required for information to be passed from one neuron to another, which is set by the time constant of the neuron (see Supplementary Information in Brown et al. (2026)^12^). Thus, the graded information is only partially transferred from neuron *i -* 1 to neuron *i* before the active gating signal advances to neuron *i* +1. In our model, the speed at which the gating signal advances (i.e., the number of response fields it traverses per unit time) is set by the speed of the eye movement and the width of a response field. Because information is only transferred while the gating signal is active, more information is transferred when the gaze falls within a response field for a longer duration, so that there is more time for the neurons to accumulate the signals they receive. Thus, more information is transferred for faster neuronal time constants (compare Figs. 2C and 3C to Supplementary Fig. 1) and larger response fields (Supplementary Fig. 2). In addition, more information may also be transferred for very small saccades in which the number of response fields traversed is small so that the loss of information at each transfer between neighboring response fields does not build up across as many transfers. Overall, higher speed movements, such as bigger saccades, lead to increasingly large information loss. The arguments above apply equally to pursuit, so that if a pursuit eye movement is too fast, similar loss of information will occur in the locally connected model.

In contrast, the broadly connected model successfully replicates the experimental findings: the graded decision information initially held in population 1 is transferred to the population 2 neurons across the saccade and is then transferred back to the population 1 neurons during the subsequent smooth pursuit eye movement (Fig. 3E,F). Information is not lost because neurons receive information directly from all neurons in the population, enabling graded evidence to be transferred directly across non-neighboring response fields. Further, the preservation of information in the circuit is independent of the time constant of the neuron and duration of the position signal (see Methods, *Model conditions for perfect integration*); these parameters affect only the rate of transfer, not whether the information will ultimately be transferred.

### Spatio-temporal dynamics of neuronal activity underlying graded evidence transfer across saccades

We further examined the spatial and temporal features of the information transfer across a saccade in the experimental data to test different assumptions of the locally and broadly connected models. First, we considered how the spatial profile of activity changed across the time course of a saccade to see if information appeared to be transferred discontinuously between neurons with distant response fields (saltatory transfer) or in a continuous, sequential sweep through neurons with adjacent response fields (fast-sweep transfer). A saltatory transfer requires broad connectivity, whereas local connectivity only has the potential to support a fast-sweep transfer. To distinguish these two possibilities, we analyzed neurons whose response fields were aligned with the choice target when the gaze was approximately midway between the fixations before and after the saccade (n= 193 neurons; see Methods). We refer to these neurons as midway neurons (Fig. 4A, right). If the graded information transfers like a fast sweep during the saccade, we expect transient activation in these neurons as the information propagates through them during the saccade. Conversely, if information transfers discontinuously, the information should bypass the midway neurons during the saccade, instead appearing only in neurons with response fields aligned to the target before and after the saccade, and appear in the midway neurons only during the pursuit.

**Figure 4.**
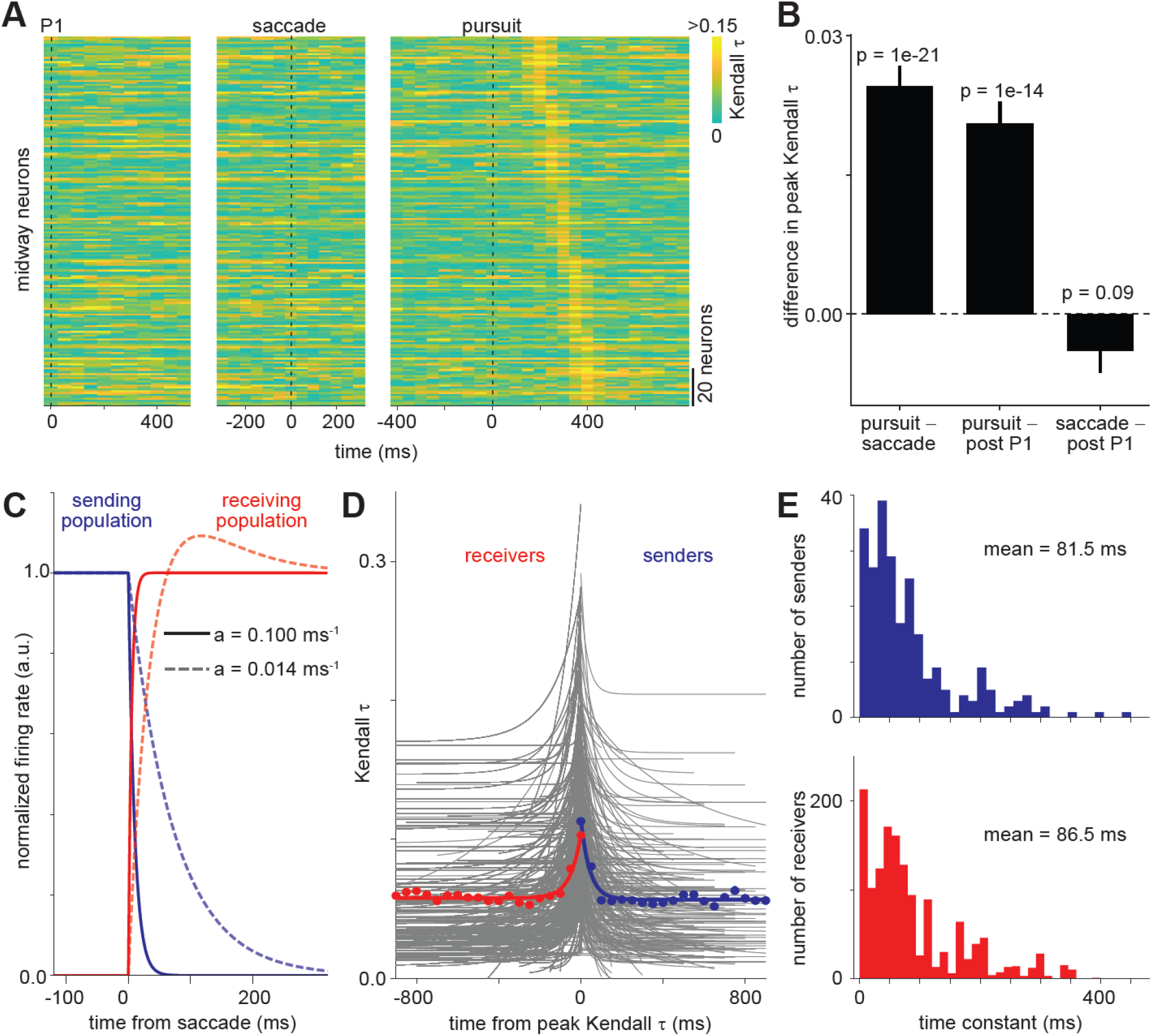
Spatio-temporal dynamics of information transfer across a saccade. (A) Heatmaps showing the decision information represented by the midway neurons (rows), aligned to the first motion pulse (left), the saccade (middle), and the pursuit (right). (B) Differences in peak information between different task epochs — *Post P1* (100–400 ms after the pulse), *Saccade* (±150 ms around saccade onset), and *Pursuit* (200–400 ms after pursuit onset) — calculated separately for each midway neuron and then averaged across neurons. Error bars denote s.e. in the population average. (C) Model simulations of the activity around the saccade, normalized to the peak of the sending population, for two different neuronal decay rates *a* (Eqs. 1, 2). Senders (blue) and receivers (red) show overlapping activity, and thus overlapping representations of decision information, regardless of the decay rate. (D) The dynamics of information content in the activity of neurons that lose (senders) and gain (receivers) information around the time of the saccade, aligned to the time of the peak information. The gray traces show the fits of the individual neuronal representations to an exponential function (Methods, Eq. 3). Note that the traces have different lengths depending on when the representation peaks for individual neurons. Bold colored lines show the average for the senders (blue) and receivers (red). (E) Histograms showing the counts of the fit time constant for the sender neurons (*top*; median = 57.1 ms) and receiver neurons (*bottom*; median = 59.6 ms). Their means (two-tailed t-test, p = 0.36) and distributions (Mann-Whitney U-test, p = 0.28; Kolmogorov-Smirnov test, p = 0.14) are not significantly different.

Midway neurons did not carry the graded information during the saccade (Fig. 4A, middle). For each midway neuron, peak decision information during the pursuit eye movement was significantly greater than during the saccade and the “Post P1” period immediately following the first motion pulse (Fig. 4B, left and middle bars; p = 1e-21, 1e-14). Peak decision information during the saccade and Post P1 did not differ significantly (Fig. 4B, right bar, p = 0.09). These results suggest that the decision information is transferred directly between neurons with distant response fields, bypassing midway neurons. This saltatory transfer favors the broadly, but not the locally, connected model.

Both the locally and broadly connected models predict that the information should decay in the sending population of neurons at the same rate that it is acquired in the receiving population (Fig. 4C), so that the total information in the system remains constant in the absence of input. Further, as described in the preceding section, the locally connected model, unlike the broadly connected model, requires a fast transfer of information between adjacent neurons to mitigate information loss. To examine these model predictions, we estimated the time constants describing the decay and acquisition of information by the populations of neurons active before and after the saccade, respectively. Our experimental data support the prediction that the neurons lose and gain information at the same rate. For this analysis, we defined “sender” and “receiver” populations of neurons as those neurons that have graded information only before or only after the saccade. For each sender and receiver neuron, we fit an exponential function (Methods, Eq. 3) to the evolution of the information contained in the neuronal firing about the motion coherence of the first pulse (i.e., the correlation between these) to get the time constant *τ*. Consistent with the models, the magnitude of the time constants of individual cells did not differ between the sending and receiving populations (Fig. 4D,E; median *τ* = 57.1 ms (senders), 59.6 ms (receivers); p = 0.36 (two-tailed t-test)). These observed timescales are far too slow for the locally connected model to faithfully transmit information between neurons during a saccade and also may be too slow even for the pursuit-only task (Supplementary Figure 1). Together, the experimental data—comprising the spatial profile of activity and the temporal scale of information transfer—are consistent with the broadly connected model, but not with the locally connected model.

## Discussion

Our broadly connected model provides an explanation for how graded information can be flexibly transferred between neurons. We propose that neurons broadcast graded information throughout a large subpopulation of neurons that may need access to this information, while a gating signal determines which neurons are receptive to the widely broadcast information. We found experimental support for this model in the observed spatio-temporal representations of graded information during a visual decision-making task. Below we discuss potential circuit and cellular mechanisms that can mediate this capability to flexibly transfer information.

### Identification of the receptive neurons

The proposed mechanism requires that the gating signal is directed to the appropriate set of neurons. These are neurons with response fields that overlap the point in retinotopic space defined by subtraction of (i) the vector defining the intervening eye movement, from (ii) the vector defining the response field of the neurons currently holding the information (i.e., the direction from the current gaze location to the choice target). As our focus is on the transfer of information, we do not explicitly model this vector subtraction, which has been treated in previous work^34–40^. Rather, we assume the existence of a circuit that achieves the vector subtraction and identifies the subpopulation for gating. This operation could be facilitated by either an efference copy triggered by the saccade itself^41–45^ or preparatory activity before the saccade^11,46^. Supporting the latter possibility, studies have shown that, when monkeys make two sequential saccades, neurons in the superior colliculus concurrently represent both saccades before the first saccade is initiated, consistent with a preparatory mechanism that precedes eye movements^47–49^.

More generally, gating signals could derive from a multitude of sources such as direct sensory inputs, attention signals^50–55^, or internal representations of spatial location. In the task modeled here, the choice target itself could provide a visual stimulus input to the cells with response fields containing it and thus be the source of the gating signal, albeit only after the eye movement. Within LIP, cells have been shown to represent spatial attention as well as integrated evidence^56^, so that spatial attention could provide the gating signal. In the locomotion-based evidence accumulation task^57^ that motivated the locally connected version of this model^12^, the gating signal could result from hippocampal or prefrontal place cell inputs^58–61^. Such candidate sources of the gating signal might be tested with perturbation experiments that inhibit a candidate brain region and monitor whether information is successfully transferred.

### Hypothesized biophysical mechanisms of the gating signal

Our rate-based models assume a simple, additive gating signal. This might be achieved mechanistically through an excitatory current received by the soma, raising the activity of the cell above its firing threshold. However, other gating mechanisms are possible, and these may be multiplicative as well as additive^62–65^. The gating signal might act presynaptically, modulating synaptic boutons (for example, through pathways mediated by G protein-coupled receptors^66^) and potentially influencing input-specific dynamics. Alternatively, the gating signal might alter dendritic receptivity, perhaps through changes in dendritic excitability^67–69^ or synaptic integration^70^. These hypotheses are neither exhaustive nor mutually exclusive; gating mechanisms may combine or vary depending on the context. Future studies using optogenetics, pharmacological manipulations, or *in vivo* imaging can help disentangle these mechanisms, providing insight into how gating enables flexible neural computations.

### Experiments to test model predictions

Our broadly connected model makes several experimentally testable predictions. First, it predicts that information transfer occurs at the same rate for both sending and receiving populations (Fig. 4C). We have shown that this is the case in the data from So & Shadlen (2022)^11^ (Fig. 4D,E).

Second, the broad connectivity predicts that the animal should be able to accommodate information transfer across saccades to any location. However, it is possible that the actual neural connections do not support every possible movement, and instead employ a system of broad connections that support typical movements. Connections between less common movements could be learned through repeated actions, thereby expanding the behaviors over which information transfer is supported. Future experiments can test this prediction by probing the behavioral performance and neural representation of the decision while the monkey makes intervening eye movements to novel locations. The presence of local versus broad connectivity could similarly be studied in navigation-based tasks in which teleportation is introduced.

### A new perspective on graded information transfer

Our work presents a new perspective on how graded information may be transferred between neurons by changing the receptivity of neurons without changing their synaptic connectivity. This approach eliminates the need to target the graded information to specific neurons. Instead, the information can be broadcast to the entire population, with specific neurons changing their receptivity due to input from a separate gating signal. This mechanism enables any neuron to receive and represent information flexibly, depending on various behavioral demands, such as different gaze directions.

In our task, the location of the choice targets remains fixed in the world (i.e., allocentrically), while shifting the gaze direction changes the oculocentric coordinates of these targets. By faithfully transferring the information between neural populations that represent the information in an oculocentric manner, the decision-related information is preserved across this coordinate change. Thus, the network is able to compensate for the shift in oculocentric reference frame in a manner that maintains integrated information about the two choices and enables the population as a whole to effectively approximate a world-centered (allocentric) frame without explicitly representing it.

While this model successfully captures much of the observed data in a task involving a single binary decision, further study is needed to understand how information is maintained and transferred in more complex tasks, including tasks in which decisions are made based on multidimensional inputs^71^. Previous studies have suggested that the transfer of graded information may not always occur between sequentially activated sets of distinct neurons, as in the current study, but instead may occur through sequentially activating different patterns of neural firing activity within the same population^72,73^. This scenario cannot be handled by the current model, in which the gating signal operates at the level of individual neurons so that distinct neurons are active before and after the transfer of information. Instead, a more complex gating mechanism would be required that allows individual neurons to remain active across the information transfer, but with different receptivity to their inputs. This might be accomplished, for example, by gating the local dendritic compartments that mediate the transfer of information between neurons^74–77^. Yet, at its essence, such a mechanism would still rely on changing receptivity to widely broadcast signals.

## Methods

### Gated receptivity model

As in Equation (1), the firing rate of a neuron in the left-preferring population of the locally connected model is given by

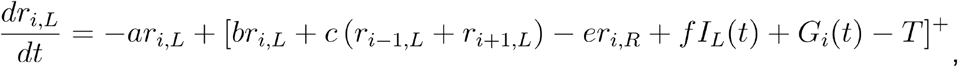

while for the broadly connected model (Eq. (2)), this firing rate is given by

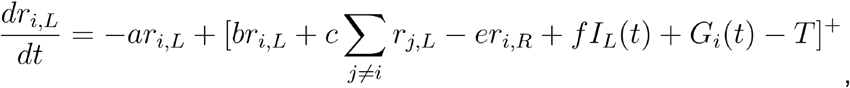

and analogous equations govern the neurons in the right-preferring population. In both models,

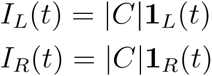

where *C* is the coherence of the random dot motion,

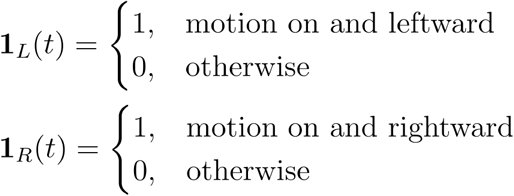

and

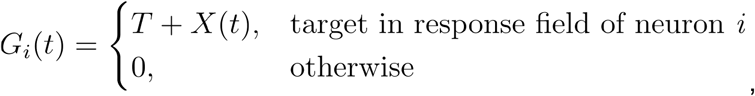

where *X*(t) represents the (potentially time varying, although held constant for simplicity in our simulations) suprathreshold component of the gating signal, and the other parameters are as described in the main text. For simulations of the locally connected model (Figs. 2C,D, 3C,D), we used parameter values *a* = 0.1 ms^-1^, *b* = 0.02 ms^-1^, *c* = 0.1 ms^-1^, *e* = 0.08 ms^-1^, *f* = 0.2 Hz/ms, *T* = 300 Hz/ms, and *X*(*t*) = 2.5 Hz/ms. For simulations of the broadly connected model (Figs. 2E,F, 3E,F) and simulations of the locally connected model at a slower timescale (Supplementary Fig. 1), we used parameter values *a* = 0.014 ms^-1^, *b* = 0.003 ms^-1^, *c* = 0.014 ms^-1^, *e* = 0.011 ms^-1^, *f* = 0.2 ms^-1^, *T* = 300 Hz/ms, and *X(t)* = 0.25 Hz/ms.

For the locally connected model in the main text, the decay rate *a* was chosen to be on a fast timescale (∼10 ms), since this model requires faster timescales to preserve information across transfers between neurons. For the broadly connected model and the locally connected model in the Supplementary Information, we use a longer time constant (∼70 ms) consistent with the typical time constants fit to the experimental data (Fig. 4D,E). Other parameters were chosen to produce firing rates in a similar range as the experimental data.

In all cases, the firing rate of the neurons at the initial gaze position were initialized to 10 Hz, and all other neuronal firing rates were initialized to 0 Hz. Note that for either model to perform integration perfectly, we require *e* = *a* - *b*. For a successful transfer between positions (given sufficient time for the transfer to occur), we require *a = c*. See the following section for a proof that, under these conditions, the broadly connected model perfectly integrates the input signals and maintains the integrated value across transfers.

### Model conditions for perfect integration

We show here that the system of differential equations defined by Equation (2) results in perfect integration of the input. Making the substitution,

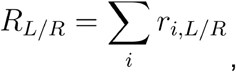

we have

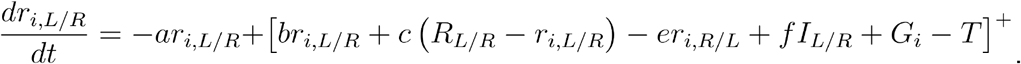

At any time only one left-right pair of units *i* =1, 2, … *N* is active. Let *k* be the active unit. For all units other than unit *k*, we have that the term in square brackets [·]^+^, and for unit *k* there is no rectification. Thus, summing equation (1) over *i* gives.

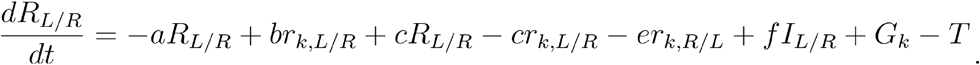

Now define *R* = *R*_*L*_−*R*_*R*_. This gives

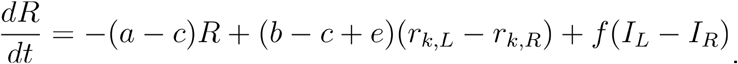

Thus, if *a - c = b - c + e = 0*,

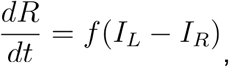

and the total activity difference integrates the input, independent of position.

### Two-pulse task

In both variants of the two-pulse task (Fig. 1A,B), animals were presented with two brief (80ms) pulses of random dot motion (P1 and P2), separated by an intervening eye movement to a choice-neutral target (T0). This choice-neutral target, shown in gray or blue, was visually distinct from the choice targets shown in red. The two pulses always conveyed the same direction of motion, but their motion strengths were independently selected from the set {0%, 4%, 8%, 16%, 32%, 64%}. There were two types of intervening eye movements, defining the two task variants. Monkeys performed each variant of the task on separate days.

### Pursuit-only variant

Animals initiated each trial by fixating at the center of the screen. While the monkey maintained central fixation, the first motion pulse (P1) was presented for 80 ms. After a variable delay of duration 700 – 1100 ms following the offset of P1, the fixation point gradually moved for 733 ms in a smooth path from the center of the screen to the gray choice-neutral target (T0), cueing the monkey to initiate and follow the fixation point with a smooth pursuit eye movement to T0. After reaching T0, the monkey maintained fixation for a variable delay of duration 200–600 ms, after which a second motion pulse (P2) was presented. Following a 500 ms delay, the fixation point at T0 disappeared, cueing a saccade to one of the two choice targets. For full task details, see So & Shadlen (2022), *Behavioral Tasks: Two-Pulse Task, 2nd Variant*^11^.

### Saccade-and-pursuit variant

This variant was similar to the pursuit-only version, with three key differences: (i) the choice-neutral target T0 was blue; (ii) after the first go cue, the monkey made a saccade to T0; and (iii) after maintaining fixation at T0 for 500–600 ms, the monkey made a smooth pursuit eye movement back to the original central fixation. As a result, the second motion pulse (P2) was viewed from the same gaze direction as the first pulse (P1). For full task details, see So & Shadlen (2022), *Behavioral Tasks: Two-Pulse Task, 1st Variant*^11^.

### Neural recordings

The main dataset comprises 832 well-isolated single neurons from LIP, recorded over 149 sessions. In each session, either a single-channel tungsten electrode (Thomas recordings; 72 sessions) or a 24-channel V-probe (Plexon Inc.; 77 sessions) was lowered through a grid to the ventral part of LIP^78^. All training, surgery, and recording procedures were in accordance with the National Institutes of Health Guide for the Care and Use of Laboratory Animals and approved by the Columbia University Institutional Animal Care and Use Committee. For complete details of the neural data collection and cell isolation, see So & Shadlen (2022)^11^.

### Cell categorization

We classified experimentally recorded neurons as belonging to population 1 or population 2 based on their response fields and decision-related activity. In sessions using single-channel electrodes, choice targets were positioned relative to the neuron’s response field so that the recorded neuron belonged to either population 1, exhibiting decision-related activity when the gaze was directed at the central fixation location, or population 2, exhibiting decision-related activity when gaze was directed away from the central fixation location. In sessions using multi-channel probes, this strategy could not be applied simultaneously to all recorded neurons, so neurons were categorized *post hoc* based on the task epoch in which they exhibited sustained decision-related activity. Decision-related activity was defined as firing rates significantly correlated with the strength and sign of motion evidence (P1 before the second motion pulse, or P1+P2 after it), measured using Kendall *τ* (p<0.05).

In the pursuit-only variant, population 1 neurons exhibited decision-related activity only during the first motion pulse (P1), whereas population 2 neurons exhibited decision-related activity during or after the intervening eye movement. In the pursuit-and-saccade variant, population 1 neurons showed decision-related activity at the central fixation, both before and after the intervening eye movements, whereas population 2 neurons showed decision-related activity when the gaze was directed away from the central fixation during the intervening eye movements. Full details of the classification procedure are described in So & Shadlen (2022)^11^.

### Analysis of the spatial organization of information transfer during saccades

To investigate how information transfers across neurons during saccades (Fig. 4A,B), we identified a subset of neurons whose response fields were aligned with the choice target when the monkey’s gaze was midway between the central fixation and the initial saccade target T0. We refer to these as *midway neurons*. To identify these neurons, we calculated the correlation, using the Kendall *τ* correlation coefficient (KC), between the motion strength in P1 and the neuron’s spike count in a 300 ms-wide bin. KC was computed in 50 ms steps from 400 ms before to 800 ms after pursuit onset. Neurons were classified as midway neurons if they had peak KC during the midpoint period (defined as 200–400 ms after pursuit onset) of the pursuit epoch and this KC was greater than 0.05. This criterion ensured that the selected neurons had response fields aligned with the choice target when the gaze was directed approximately midway between the central fixation and T0. For these midway neurons, we compared their decision representation during the Post P1 epoch (100–400 ms after the pulse), the peri-Saccade epoch (-150–150 ms around the saccade onset), and the Pursuit epoch (200–400 ms after pursuit onset).

### Analysis of temporal dynamics of information transfer during saccades

For the analysis of the temporal dynamics of information transfer (Fig. 4D,E), we considered transfer around the time of the intervening saccade in the saccade-and-pursuit variant of the task. To identify neurons sending and receiving the decision information around the intervening saccade, we calculated the correlation, using the Kendall *τ* correlation coefficient (KC), between the motion strength in P1 and the neuron’s spike count in 300 ms-wide bins before (-450 – -150 ms) and after (150 – 450 ms) the intervening saccade. Senders are defined as those neurons with correlation coefficients bigger than 0.05 before the saccade and not bigger than 0.05 after the saccade. Receivers are neurons with coefficients bigger than 0.05 after the saccade and not bigger than 0.05 before. We identified 245 sender and 1438 receiver neurons.

To estimate how information about the motion strength appeared and disappeared in the sending and receiving neurons, we first calculated the time at which each neuron exhibited its peak correlation. Then, for the sending population, we calculated how this information decayed over subsequent time bins after the peak. For the receiving population, we calculated how the information grew over preceding time bins to its peak. To do this, we recalculated KC in a sliding window of 100 ms that advanced in 50 ms steps and fit the resulting values to an exponential function of the form for each neuron

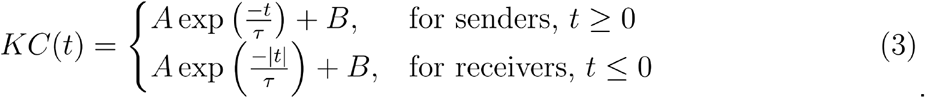

Differences in the means of the fit time constants *τ* from the sending and receiving populations were tested for significance using a two-tailed t-test, and differences in the distributions were tested for significance using a Mann-Whitney U-test and a Kolmogorov-Smirnov test (Fig. 4D,E). For statistical comparisons, outliers were excluded separately for sender and receiver populations using the interquartile range (IQR) method (excluding datapoints more than 1.5 × IQR from the boundaries of the interquartile range). Results were qualitatively unchanged when analyses were performed without excluding outliers (p=0.86 for t-test; p=0.57 for Mann-Whitney U-test; p=0.27 for Kolmogorov-Smirnov test).

### Model simulations

We simulated the models on both the pursuit-only (Fig. 2) and saccade-and-pursuit variant (Fig. 3) using scipy’s odeint function, which uses the lsoda algorithm. We used a maximum step size of 0.1 ms to find the activity levels of the neurons in each task in 0.1 ms increments in the trial.

In the simulations in Figs. 2 and 3 and Supplementary Fig. 1 and 3, we used *N* = 10 neurons in each of the left-and right-preferring populations. In Supplementary Fig. 2, we used *N* = 5. We defined neuron 0 in each population as the population 1 neuron and all other neurons as population 2 neurons. We defined neuron *N* - 1 as the neuron with gaze direction corresponding to the target T0. We assumed the response field of the other neurons in the population were distributed evenly and sequentially between these two neurons.

Simulations were performed assuming constant inputs during the motion pulses. Simulations were allowed to equilibrate for 100 ms before initiating the trial (times from -100 - 0 ms). The timings of the start and end of the intervening eye movements were drawn uniformly from within a 50 ms window to reflect variance in reaction times and movement speeds across trials. Tables 1-4 summarize the values of the different functions at different time points in the simulation.

**Table 1.**
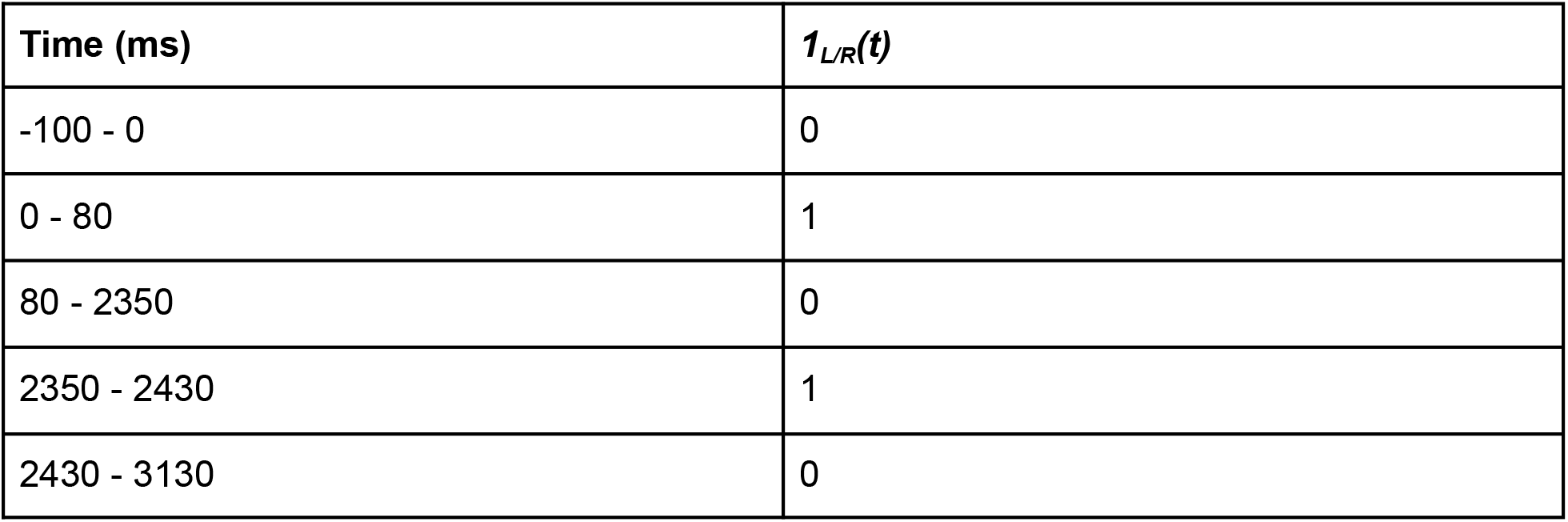
Values of the random dot motion indicator function during the pursuit-only variant. For motion coherence C ≥ 0, this table gives values for ***1***_***L***_(*t*) while ***1***_***R***_(*t*) = 0 for all *t*. For *C* ≤ 0, ***1***_***R***_(*t*) takes the shown values while ***1***_***L***_(*t*) = 0 for all *t*.

**Table 2.** Values of the gating signal during the pursuit only variant. Here, pursuit_start_ = 980 ms + ε, pursuit_end_ = 1790 ms + δ, and ε and δ are random uniform variables in the range (-25, 25) ms, drawn independently for each trial.

| Time (ms) | $\mathbf{G}_i(t)$ |
| --- | --- |
| -100 - pursuit<sub>start</sub> | $T+X(t)$, for $i = 0$<br>0, for $i \neq 0$ |
| pursuit<sub>start</sub> - pursuit<sub>end</sub> | $T + X(t)$, for $i \leq \frac{N(t - \text{pursuit}_{\text{start}})}{(\text{pursuit}_{\text{end}} - \text{pursuit}_{\text{start}})} < (i + 1)$<br>0, otherwise |
| pursuit<sub>end</sub> - 3130 | $T+X(t)$, for $i = (N-1)$<br>0, for $i \neq (N-1)$ |

**Table 3.** Values of the random dot motion indicator function during the saccade-and-pursuit variant. For motion coherence C ≥ 0, this table gives values for ***1***_***L***_(*t*) while ***1***_***R***_(*t*) = 0 for all *t*. For *C* ≤ 0, ***1***_***R***_(*t*) takes the shown values while ***1***_***L***_(*t*) = 0 for all *t*.

| Time (ms) | $1_{L/R}(t)$ |
| --- | --- |
| -100 - 0 | 0 |
| 0 - 80 | 1 |
| 80 - 2900 | 0 |
| 2900 - 2980 | 1 |
| 2980 - 3680 | 0 |

**Table 4.** Values of the gating signal during the saccade-and-pursuit variant. Here saccade_start_ = 680 ms + ε, saccade_end_ = saccade_start_ + 10 ms, pursuit_start_ = 1450 ms + δ, pursuit_end_ = 2200 ms + γ, and ε, δ, and γ are random uniform variables in the range (-25, 25) ms, drawn independently for each trial. Note that assuming a longer, 20 ms saccade duration does not rescue the graded information transfer in the locally connected model (Supplementary Fig. 3).

| Time (ms) | $G_i(t)$ |
| --- | --- |
| -100 - saccade<sub>start</sub> | $T+X(t)$, for $i = 0$<br>0, for $i \neq 0$ |
| saccade<sub>start</sub> - saccade<sub>end</sub> | In simulations of the broadly connected model (assuming a step change in gaze direction):<br>$T+X(t)$, for $i = (N-1)$<br>0, for $i \neq (N-1)$<br>In simulations of the locally connected model (allowing a fast sweep of gaze direction):<br>$T + X(t)$, for $i \leq \frac{N(t - \text{saccade}_{\text{start}})}{(\text{saccade}_{\text{end}} - \text{saccade}_{\text{start}})} < (i + 1)$<br>0, otherwise |
| saccade<sub>end</sub> - pursuit<sub>start</sub> | $T+X(t)$, for $i = (N-1)$<br>0, for $i \neq (N-1)$ |
| pursuit<sub>start</sub> - pursuit<sub>end</sub> | $T + X(t)$, for $((N - 1) - i) \leq \frac{N(t - \text{pursuit}_{\text{start}})}{(\text{pursuit}_{\text{end}} - \text{pursuit}_{\text{start}})} < (N - i)$<br>0, otherwise |
| pursuit<sub>end</sub> - 3130 | $T+X(t)$, for $i = 0$<br>0, for $i \neq 0$ |

## Data Availability

All neural data has been previously collected and published^11^. It is available at: https://data.mendeley.com/datasets/ptcvtxg55j/1

## Code Availability

All code for the models and data analysis is available at: https://github.com/lindseysbrown/gated_graded_information_transfer/

## Acknowledgements

We thank Gregory Davis for helpful discussions about this work. Funding was provided by NIH F32MH132179 (LSB), NIH R01NS113113 (MNS), NIH R01MH122513 (MNS), NIH U19NS132720 (MSG), a Burroughs Wellcome Fund Career Award at the Scientific Interface (LSB), the Sloan Swartz Foundation (LSB), the Howard Hughes Medical Institute (HHMI; NS, MNS), the Grossman Center for Statistics of the Mind (MNS), the Air Force Office for Scientific Research 21RT0878 (MNS), and the Kavli Center, Columbia University (MNS).

## Supplementary Information

**Supplementary Figure 1.**
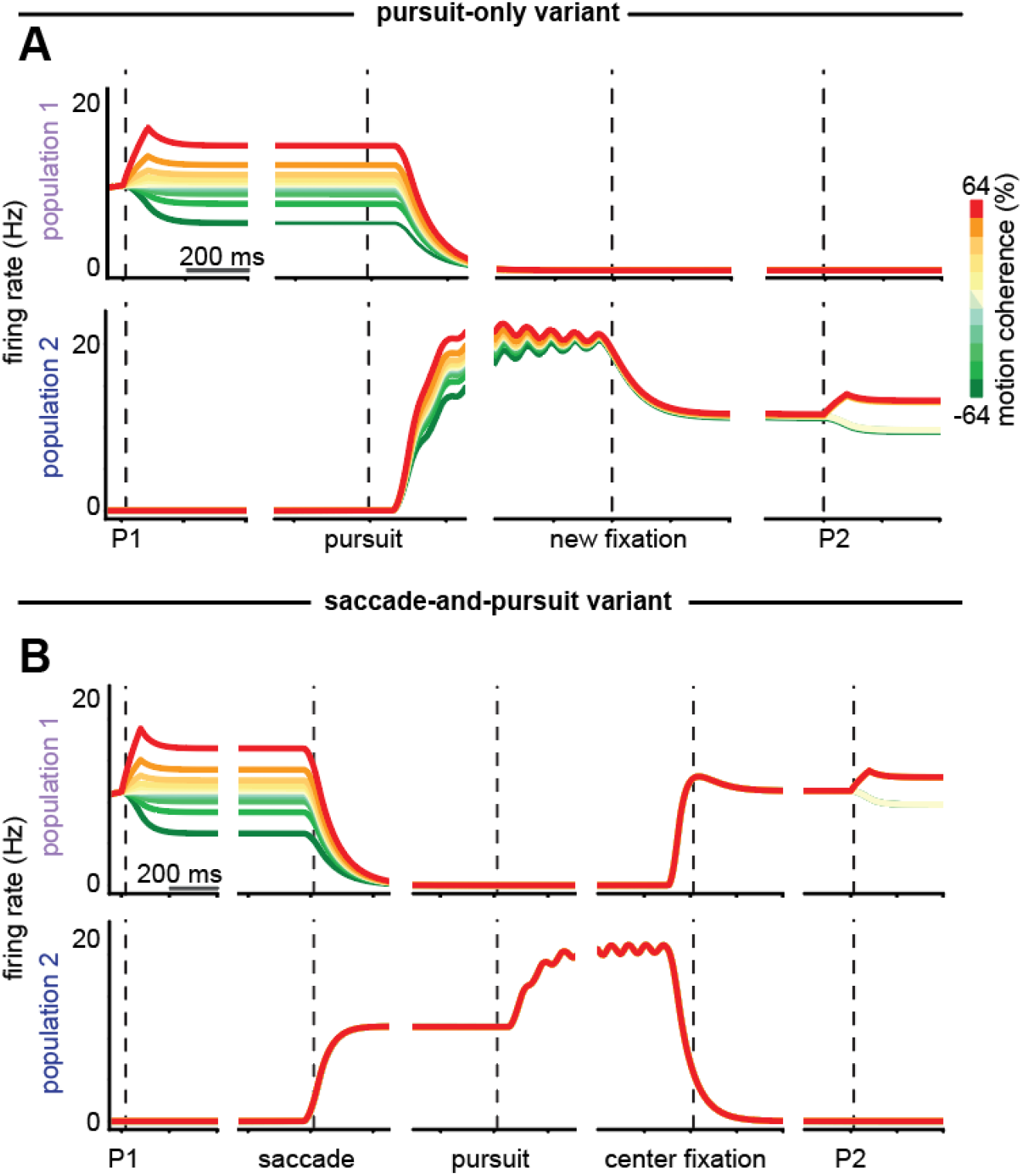
Slower neuronal time constants decrease information transfer in the locally connected model. (A) Sum total of the trial-averaged activity of the simulated neurons in each population during the pursuit-only variant (*top*: population 1, *bottom*: population 2). Colors indicate motion strength and direction. (B) Same as A, but for the saccade-and-pursuit variant. Compared to the simulations in Figs. 2 and 3, where we use a relatively fast time constant (*a* = 0.1 ms^-1^), here we use a longer time constant (*a* = 0.014 ms^-1^) more similar to those typically observed in the experimental data (Fig. 4). With the longer time constant, more information is lost at each transition between gaze directions (note the decrease in the spread of the graded information during the pursuit in population 2 in A). Thus, the locally connected model fails to transfer graded information even in the pursuit-only case for longer time constants, unlike the broadly connected model which is robust to these longer time constants for both variants of the task (Figs. 2E, 3E).

**Supplementary Figure 2.**
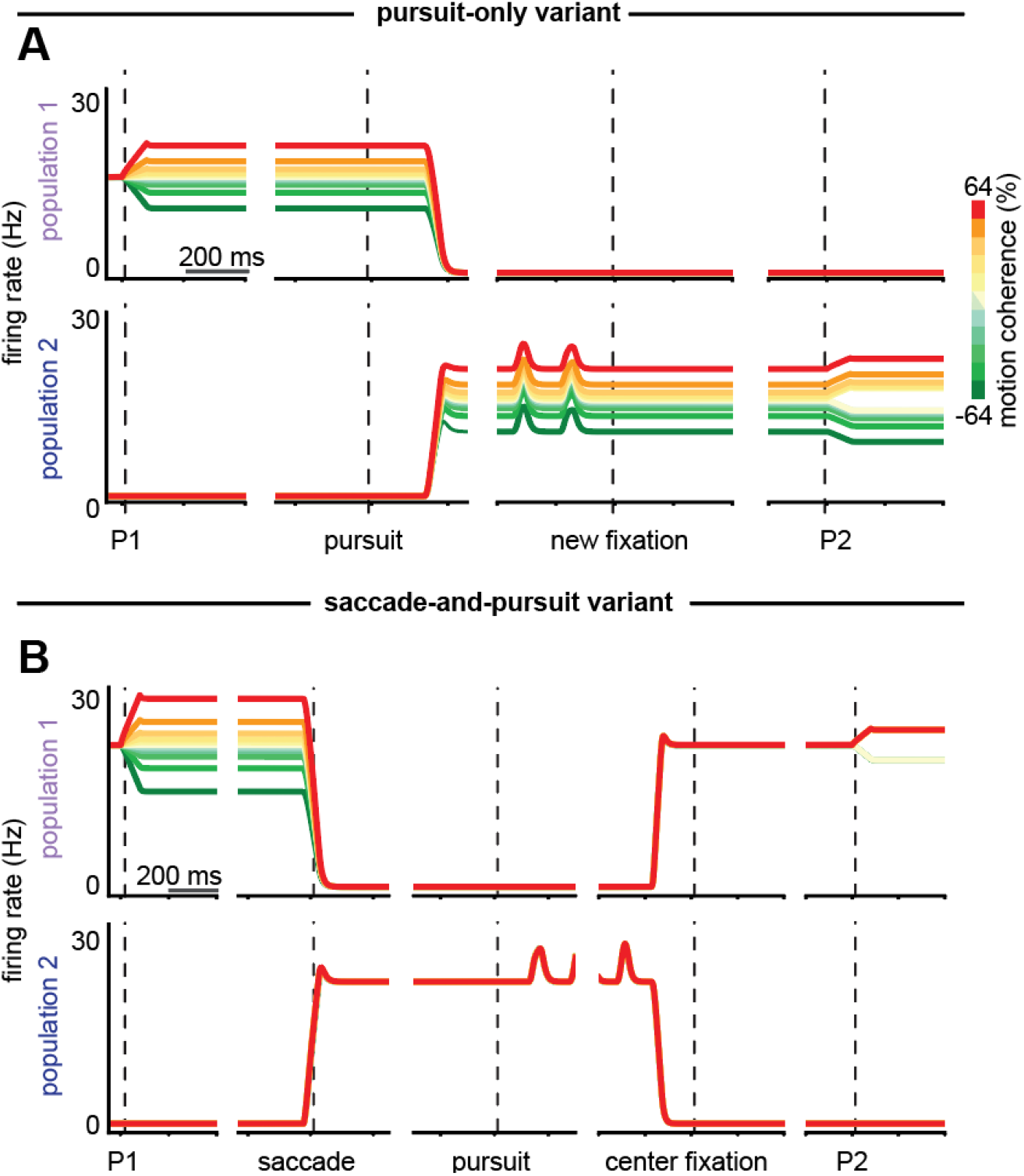
Wider response fields do not rescue graded evidence transfer across saccades in the locally connected model. (A) Sum total of the trial-averaged activity of the simulated neurons in each population during the pursuit-only variant (*top*: population 1, *bottom*: population 2). Colors indicate motion strength and direction. (B) Same as A, but for the saccade-and-pursuit variant. Compared to the simulations in Figs. 2-3, where we assume the gaze direction traverses a large number of response fields (10 neurons in each of the left- and right-preferring populations; alternatively thought of as a longer saccade), here we use wider response fields (5 neurons in each of the left- and right-preferring populations) more similar to the reported widths of response fields for the relatively short saccade used in this experiment. Thus, the limitations we observed in the locally connected model do not result from our relatively small choice of response field size. However, if response fields were so large (or the saccade so small) that the saccade passed through only one transition, i.e. did not need to be transferred through any intermediate response fields, then the graded evidence could be transferred through a single local connection. In contrast, the broadly connected model is robust to both the size of the response fields and the length of the saccade.

**Supplementary Figure 3.**
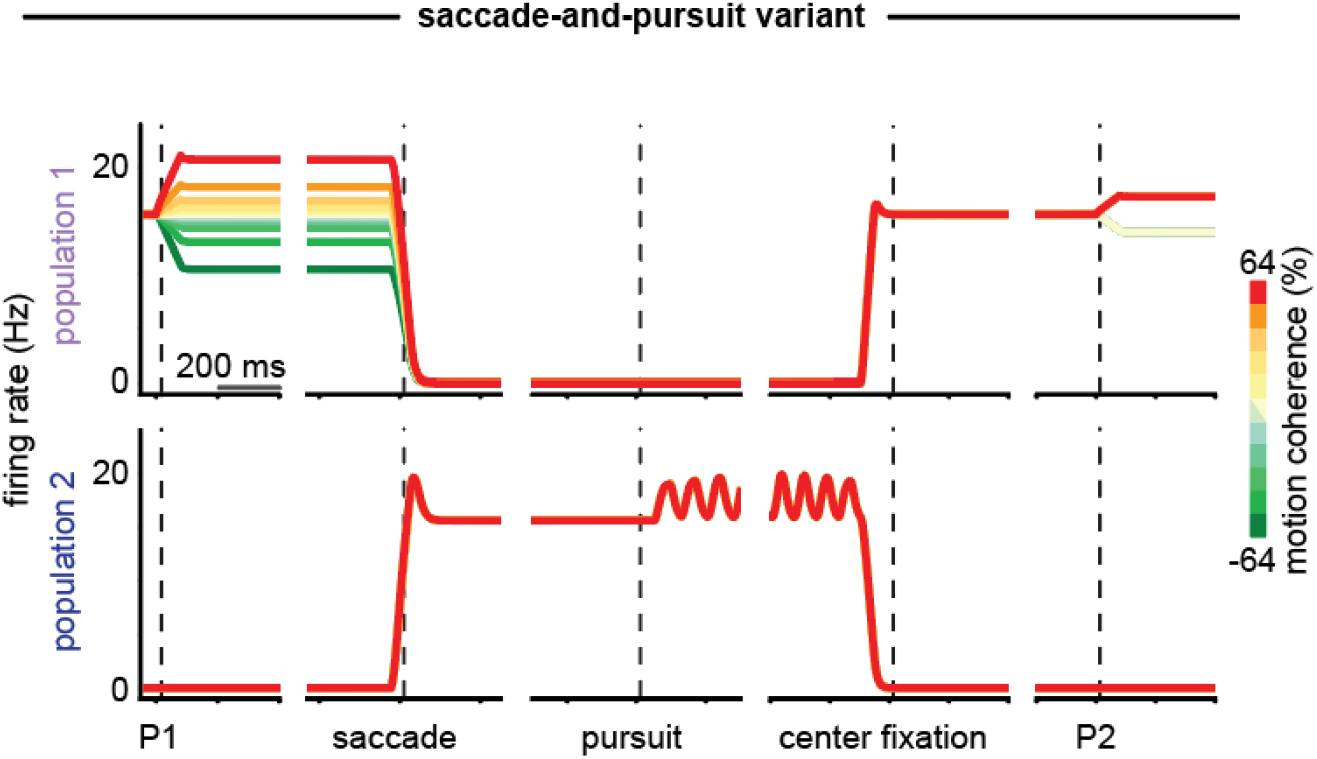
Slower saccades do not rescue graded evidence transfer across saccades in the locally connected model. Sum total of the trial-averaged activity of the simulated neurons in each population during the saccade-and-pursuit variant (*top*: population 1, *bottom*: population 2). Colors indicate motion strength and direction. Compared to the simulations in Fig. 3, where we assume the saccade lasts 10 ms, here we assume a slower saccade with duration 20 ms. Despite the increased duration of the saccade, the gating signal does not dwell long enough within each receptive field to transfer a graded representation across the saccade.

## References

1. Shadlen, M. N. & Newsome, W. T. Neural basis of a perceptual decision in the parietal cortex (area LIP) of the rhesus monkey. J. Neurophysiol. 86, 1916–1936xs (2001).

2. Cisek, P. & Kalaska, J. F. Neural mechanisms for interacting with a world full of action choices. Annu. Rev. Neurosci. 33, 269–298 (2010).

3. Lindner, A., Iyer, A., Kagan, I. & Andersen, R. A. Human posterior parietal cortex plans where to reach and what to avoid. J. Neurosci. 30, 11715–11725 (2010).

4. Shadlen, M. N. & Newsome, W. T. Motion perception: seeing and deciding. Proc. Natl. Acad. Sci. U. S. A. 93, 628–633 (1996).

5. Shadlen, M. N., Kandel, E. R. Decision-making and consciousness. in Principles of Neural Science, 6e (ed. E.R. Kandel, J.D. Koester, S.H. Mack, S.A. Siegelbaum) (McGraw Hill, 2021).

6. Harvey, C. D., Coen, P. & Tank, D. W. Choice-specific sequences in parietal cortex during a virtual-navigation decision task. Nature 484, 62–68 (2012).

7. Koay, S. A., Charles, A. S., Thiberge, S. Y., Brody, C. D. & Tank, D. W. Sequential and efficient neural-population coding of complex task information. Neuron 110, 328–349.e11 (2022).

8. Krumin, M., Lee, J. J., Harris, K. D. & Carandini, M. Decision and navigation in mouse parietal cortex. Elife 7, (2018).

9. Terada, S., Sakurai, Y., Nakahara, H. & Fujisawa, S. Temporal and Rate Coding for Discrete Event Sequences in the Hippocampus. Neuron 94, 1248–1262.e4 (2017).

10. Nieh, E. H. et al. Geometry of abstract learned knowledge in the hippocampus. Nature 595, 80–84 (2021).

11. So, N. & Shadlen, M. N. Decision formation in parietal cortex transcends a fixed frame of reference. Neuron 110, 3206–3215.e5 (2022).

12. Brown, L. S. et al. Neural circuit models for evidence accumulation through choice-selective sequences. Nat. Commun. (2026) doi:10.1038/s41467-026-70267-9.

13. Ditterich, J., Mazurek, M. E. & Shadlen, M. N. Microstimulation of visual cortex affects the speed of perceptual decisions. Nat. Neurosci. 6, 891–898 (2003).

14. Cook, E. P. & Maunsell, J. H. R. Dynamics of neuronal responses in macaque MT and VIP during motion detection. Nat. Neurosci. 5, 985–994 (2002).

15. Chandrasekaran, C., Peixoto, D., Newsome, W. T. & Shenoy, K. V. Laminar differences in decision-related neural activity in dorsal premotor cortex. Nat. Commun. 8, 614 (2017).

16. Peixoto, D. et al. Decoding and perturbing decision states in real time. Nature 591, 604–609 (2021).

17. Hanks, T. D. et al. Distinct relationships of parietal and prefrontal cortices to evidence accumulation. Nature 520, 220–223 (2015).

18. Bogacz, R., Hu, P. T., Holmes, P. J. & Cohen, J. D. Do humans produce the speed-accuracy trade-off that maximizes reward rate? Q. J. Exp. Psychol. (Hove) 63, 863–891 (2010).

19. Laming, D. R. J. Information Theory of Choice-Reaction Times. (Academic Press, 1968).

20. Ratcliff, R. A theory of memory retrieval. Psychol. Rev. 85, 59–108 (1978).

21. Stone, M. Models for choice-reaction time. Psychometrika 25, 251–260 (1960).

22. Bogacz, R., Brown, E., Moehlis, J., Holmes, P. & Cohen, J. D. The physics of optimal decision making: a formal analysis of models of performance in two-alternative forced-choice tasks. Psychol. Rev. 113, 700–765 (2006).

23. Usher, M. & McClelland, J. L. The time course of perceptual choice: the leaky, competing accumulator model. Psychol. Rev. 108, 550–592 (2001).

24. Goldman, M. S., Compte, A. & Wang, X.-J. Neural integrator models. Encyclopedia of neuroscience 165–178 (2009).

25. Machens, C. K., Romo, R. & Brody, C. D. Flexible control of mutual inhibition: a neural model of two-interval discrimination. Science 307, 1121–1124 (2005).

26. Lam, N. H. et al. Effects of Altered Excitation-Inhibition Balance on Decision Making in a Cortical Circuit Model. J. Neurosci. 42, 1035–1053 (2022).

27. Roitman, J. D. & Shadlen, M. N. Response of neurons in the lateral intraparietal area during a combined visual discrimination reaction time task. J. Neurosci. 22, 9475–9489 (2002).

28. Gold, J. I. & Shadlen, M. N. The neural basis of decision making. Annu. Rev. Neurosci. 30, 535–574 (2007).

29. Churchland, A. K. et al. Variance as a signature of neural computations during decision making. Neuron 69, 818–831 (2011).

30. Steinemann, N. A. et al. Direct observation of the neural computations underlying a single decision. (2024) doi:10.7554/elife.90859.2.

31. Wang, X.-J. Probabilistic decision making by slow reverberation in cortical circuits. Neuron 36, 955–968 (2002).

32. Lo, C.-C. & Wang, X.-J. Cortico–basal ganglia circuit mechanism for a decision threshold in reaction time tasks. Nat. Neurosci. 9, 956–963 (2006).

33. Wong, K.-F. & Wang, X.-J. A recurrent network mechanism of time integration in perceptual decisions. J. Neurosci. 26, 1314–1328 (2006).

34. Salinas, E. & Abbott, L. F. Decoding vectorial information from firing rates. in The Neurobiology of Computation 299–304 (Springer US, Boston, MA, 1995).

35. Pouget, A. & Sejnowski, T. J. Spatial transformations in the parietal cortex using basis functions. J. Cogn. Neurosci. 9, 222–237 (1997).

36. Quaia, C., Optican, L. M. & Goldberg, M. E. The maintenance of spatial accuracy by the perisaccadic remapping of visual receptive fields. Neural Netw. 11, 1229–1240 (1998).

37. Perrone, J. A. & Krauzlis, R. J. Vector subtraction using visual and extraretinal motion signals: a new look at efference copy and corollary discharge theories. J. Vis. 8, 24.1–14 (2008).

38. Lyu, C., Abbott, L. F. & Maimon, G. Building an allocentric travelling direction signal via vector computation. Nature 601, 92–97 (2022).

39. Groh, J. M. & Sparks, D. L. Two models for transforming auditory signals from head-centered to eye-centered coordinates. Biol. Cybern. 67, 291–302 (1992).

40. Cassanello, C. R. & Ferrera, V. P. Visual remapping by vector subtraction: analysis of multiplicative gain field models. Neural Comput. 19, 2353–2386 (2007).

41. Sommer, M. A. & Wurtz, R. H. A pathway in primate brain for internal monitoring of movements. Science 296, 1480–1482 (2002).

42. Sommer, M. A. & Wurtz, R. H. What the brain stem tells the frontal cortex. I. Oculomotor signals sent from superior colliculus to frontal eye field via mediodorsal thalamus. J. Neurophysiol. 91, 1381–1402 (2004).

43. Sommer, M. A. & Wurtz, R. H. What the brain stem tells the frontal cortex. II. Role of the SC-MD-FEF pathway in corollary discharge. J. Neurophysiol. 91, 1403–1423 (2004).

44. Bridgeman, B. A review of the role of efference copy in sensory and oculomotor control systems. Ann. Biomed. Eng. 23, 409–422 (1995).

45. Keith, G. P., Blohm, G. & Crawford, J. D. Influence of saccade efference copy on the spatiotemporal properties of remapping: a neural network study. J. Neurophysiol. 103, 117–139 (2010).

46. Paré, M. & Wurtz, R. H. Progression in neuronal processing for saccadic eye movements from parietal cortex area lip to superior colliculus. J. Neurophysiol. 85, 2545–2562 (2001).

47. Sparks, D. L. & Porter, J. D. Spatial localization of saccade targets. II. Activity of superior colliculus neurons preceding compensatory saccades. J. Neurophysiol. 49, 64–74 (1983).

48. McPeek, R. M. & Keller, E. L. Superior colliculus activity related to concurrent processing of saccade goals in a visual search task. J. Neurophysiol. 87, 1805–1815 (2002).

49. McPeek, R. M., Han, J. H. & Keller, E. L. Competition between saccade goals in the superior colliculus produces saccade curvature. J. Neurophysiol. 89, 2577–2590 (2003).

50. Moore, T., Armstrong, K. M. & Fallah, M. Visuomotor origins of covert spatial attention. Neuron 40, 671–683 (2003).

51. Sperling, G. & Melchner, M. J. The attention operating characteristic: examples from visual search. Science 202, 315–318 (1978).

52. Corbetta, M. et al. A common network of functional areas for attention and eye movements. Neuron 21, 761–773 (1998).

53. Hoffman, J. E. & Subramaniam, B. The role of visual attention in saccadic eye movements. Percept. Psychophys. 57, 787–795 (1995).

54. Kustov, A. A. & Robinson, D. L. Shared neural control of attentional shifts and eye movements. Nature 384, 74–77 (1996).

55. Nobre, A. C., Gitelman, D. R., Dias, E. C. & Mesulam, M. M. Covert visual spatial orienting and saccades: overlapping neural systems. Neuroimage 11, 210–216 (2000).

56. Seideman, J. A., Stanford, T. R. & Salinas, E. A conflict between spatial selection and evidence accumulation in area LIP. Nat. Commun. 13, 4463 (2022).

57. Pinto, L. et al. An Accumulation-of-Evidence Task Using Visual Pulses for Mice Navigating in Virtual Reality. Front. Behav. Neurosci. 12, 36 (2018).

58. Moser, M.-B., Rowland, D. C. & Moser, E. I. Place cells, grid cells, and memory. Cold Spring Harb. Perspect. Biol. 7, a021808 (2015).

59. O’keefe, J. & Nadel, L. The Hippocampus as a Cognitive Map. (Oxford University Press, 1978).

60. Sauer, J.-F., Folschweiller, S. & Bartos, M. Topographically organized representation of space and context in the medial prefrontal cortex. Proc. Natl. Acad. Sci. U. S. A. 119, e2117300119 (2022).

61. Bota, A. et al. Shared and unique properties of place cells in anterior cingulate cortex and hippocampus. bioRxiv (2021) doi:10.1101/2021.03.29.437441.

62. Jadi, M. P., Behabadi, B. F., Poleg-Polsky, A., Schiller, J. & Mel, B. W. An augmented two-layer model captures nonlinear analog spatial integration effects in pyramidal neuron dendrites. Proc. IEEE Inst. Electr. Electron. Eng. 102, 782–798 (2014).

63. Behabadi, B. F., Polsky, A., Jadi, M., Schiller, J. & Mel, B. W. Location-dependent excitatory synaptic interactions in pyramidal neuron dendrites. PLoS Comput. Biol. 8, e1002599 (2012).

64. Mehaffey, W. H., Doiron, B., Maler, L. & Turner, R. W. Deterministic multiplicative gain control with active dendrites. J. Neurosci. 25, 9968–9977 (2005).

65. Bicknell, B. A. & Häusser, M. A synaptic learning rule for exploiting nonlinear dendritic computation. Neuron 109, 4001–4017.e10 (2021).

66. Betke, K. M., Wells, C. A. & Hamm, H. E. GPCR mediated regulation of synaptic transmission. Prog. Neurobiol. 96, 304–321 (2012).

67. Dembrow, N. C. & Spain, W. J. Input rate encoding and gain control in dendrites of neocortical pyramidal neurons. Cell Rep. 38, 110382 (2022).

68. Behabadi, B. F. & Mel, B. W. Mechanisms underlying subunit independence in pyramidal neuron dendrites. Proc. Natl. Acad. Sci. U. S. A. 111, 498–503 (2014).

69. Schiller, J., Major, G., Koester, H. J. & Schiller, Y. NMDA spikes in basal dendrites of cortical pyramidal neurons. Nature 404, 285–289 (2000).

70. Mel, B. W. Synaptic integration in an excitable dendritic tree. J. Neurophysiol. 70, 1086–1101 (1993).

71. Mante, V., Sussillo, D., Shenoy, K. V. & Newsome, W. T. Context-dependent computation by recurrent dynamics in prefrontal cortex. Nature 503, 78–84 (2013).

72. Elsayed, G. F., Lara, A. H., Kaufman, M. T., Churchland, M. M. & Cunningham, J. P. Reorganization between preparatory and movement population responses in motor cortex. Nat. Commun. 7, 13239 (2016).

73. Tang, C., Herikstad, R., Parthasarathy, A., Libedinsky, C. & Yen, S.-C. Minimally dependent activity subspaces for working memory and motor preparation in the lateral prefrontal cortex. Elife 9, (2020).

74. Araya, R., Vogels, T. P. & Yuste, R. Activity-dependent dendritic spine neck changes are correlated with synaptic strength. Proc. Natl. Acad. Sci. U. S. A. 111, E2895–E2904 (2014).

75. Yuste, R. & Denk, W. Dendritic spines as basic functional units of neuronal integration. Nature 375, 682–684 (1995).

76. Harris, K. M. & Kater, S. B. Dendritic spines: cellular specializations imparting both stability and flexibility to synaptic function. Annu. Rev. Neurosci. 17, 341–371 (1994).

77. Yuste, R. Dendritic spines and distributed circuits. Neuron 71, 772–781 (2011).

78. Lewis, J. W. & Van Essen, D. C. Corticocortical connections of visual, sensorimotor, and multimodal processing areas in the parietal lobe of the macaque monkey. J. Comp. Neurol. 428, 112–137 (2000).

